# Spontaneous behavior predicts colony membership, individual identity, and rank in naked mole-rat societies

**DOI:** 10.64898/2026.09.27.754730

**Authors:** Yuki Haba, Ryan Schwark, Caleb Weinreb, Simon Ogundare, William Foster, Yu-Young Wesley Tsai, Jerry Lyu, Mayssam Mohamed, Phalaen Chang, Amanda Arnold, Evan Schaffer, Kanaka Rajan, Sandeep Robert Datta, Ishmail Abdus-Saboor

**Affiliations:** Zuckerman Mind Brain Behavior Institute, Columbia University; Department of Biological Sciences and Howard Hughes Medical Institute, Columbia University; Department of Neurobiology, Harvard Medical School; Nash Family Department of Neuroscience and Friedman Brain Institute, Icahn School of Medicine at Mount Sinai; Kempner Institute for the Study of Natural and Artificial Intelligence, Harvard University

## Abstract

Animal societies are organized across scales, from moment-to-moment actions of individuals to dynamic group structure. A central challenge in sociobiology is to understand how behaviors of individuals shape group structure. Here, using machine vision to map recurrent fine-scale behavioral modules in the eusocial naked mole-rat (*Heterocephalus glaber*), we show that spontaneous behavior in isolation predicts group-level social organization, including colony membership and individual identity. We determined dominance relationships and found highly linear hierarchies, with ranks stable over months to years. Remarkably, fine-scale behavioral profiles in isolation correctly predicted pairwise dominance relationships, allowing reconstruction of rank order within each colony. Models trained within one colony predicted rank poorly in others, suggesting that naked mole-rats express social position through behavioral patterns specific to each colony. Together, our results connect fine-scale behavior to social structure maintained over years and show that, even when an animal is alone, its behavior carries signatures of its identity, colony, and place within that society.

## INTRODUCTION

Animal societies are organized across temporal and organizational scales. An aggressive bite, vocal exchange, or affiliative contact unfolds in seconds; repeated encounters across minutes to hours form an interaction network; and over days, group membership, social bonds, and dominance relationships emerge and may endure for years. Understanding how moment-to-moment actions of individuals relate to persistent group structure is a central goal in sociobiology (1, 2).

Resolving the relationship between individual and group requires measuring behavior with temporal precision to capture individual behaviors while simultaneously determining the social organization in which they are embedded. Conventional studies have therefore faced a trade-off between behavioral resolution and social scale: deeply annotating a few interactions or coarsely following many individuals. Fortunately, recent advances in computational ethology are beginning to relax this trade-off (3). Deep learning-based pose estimators now track user-defined anatomical landmarks without physical markers across large video datasets (4, 5), while unsupervised computational methods use temporal structure in images or pose dynamics to segment continuous movement into recurring fine-scale modules, often termed “behavioral syllables,” without predefined action labels (6–8).

These computational ethology approaches are flexible across species and experimental settings, allowing researchers to connect transient movements to more persistent biological phenomena (3). For example, high-resolution analyses of behavior revealed hierarchical structure in fly behavior (9), mapped pharmacological perturbations in mice (10), and predicted aging trajectories in killifish (11). Efforts to link individual behavior to social organization remain limited, albeit promising. A handful of studies have linked specific individual traits, such as exploration or sociability, to particular social roles and collective outcomes (12–15). In mammalian societies, however, whether and how individual behaviors reflect enduring social organization remains largely unknown.

The African naked mole-rat (*Heterocephalus glaber*) offers an ideal mammalian model for studying social behavior across scales. Naked mole-rats live in large, multigenerational colonies with pronounced reproductive division of labor and dominance relationships (16, 17). Each colony consists of a single breeding queen, one to three breeding males, and dozens of non-reproductive workers that share communal tasks such as foraging, nest building, and care of offspring, forming a highly coordinated society. Naked mole-rats show robust colony recognition (18, 19) and their vocalizations carry individual-specific signatures (19). Captive colonies retain key features of this social organization, allowing researchers to investigate sociality across scales from molecules to behaviors to colony-wide dynamics (19–21). Here, we leverage high-resolution pose tracking and behavioral segmentation to show that spontaneous behavior in isolation predicts individual identity, colony membership, and position within stable dominance hierarchies in naked mole-rats.

## RESULTS

### Naked mole-rats express diverse spontaneous behavioral syllables

To characterize behavioral syllables in the spontaneous behavior of naked mole-rats, we recorded individual animals from four colonies in a circular arena with digging material (Fig. 1A; 85 animals in total, 10 minutes each). We trained a SLEAP model (4) to estimate the positions of seven body landmarks. We then used the resulting pose trajectories across 3.45 million video frames to model candidate behavioral syllables with Keypoint-MoSeq (8). We selected the temporal scale parameter (kappa) by comparing syllable-duration distributions and representative movements across a range of values, and then chose the final model among 30 randomly initialized fits by marginal likelihood (see Methods). The final model, after visual curation, identified 51 high-confidence behavioral syllables, comprising 50,728 instances with a median duration of 767 ms (Fig. 1A; fig. S1A). We grouped these syllables into seven broad syllable families: forward movement, reverse movement, pause, scratch, dig, turn, and stand up (Fig. 1A; Supplementary Video 1). Of note, backward locomotion is rare among mammals, but naked mole-rats readily travel both forward and backward through their tunnels, a behavior captured in our dataset (22).

**Figure 1.**
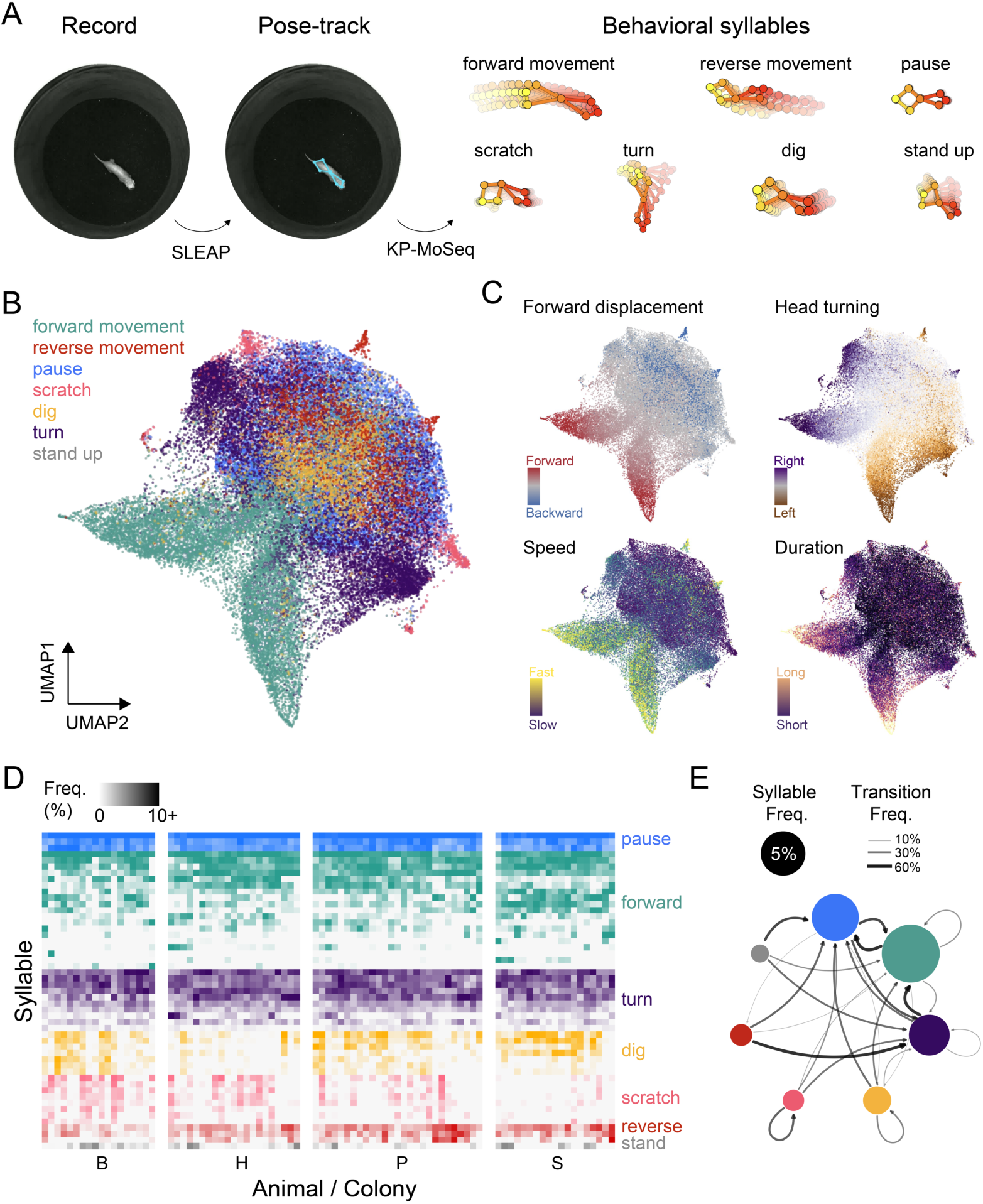
Naked mole-rats express diverse spontaneous behavioral syllables. (**A**) Behavioral syllable modeling workflow. Pose trajectories were inferred from 10-minute single-animal recordings and used to identify behavioral syllables. We identified 51 behavioral syllables and grouped them into seven syllable families based on kinematic features and visual inspection of representative instances. (**B** and **C**) UMAP embedding of 50,728 syllable instances identified across all recordings, colored by syllable family (B) or four movement features (C). (**D**) Relative syllable expression frequencies across animals. Rows and columns indicate individual syllables and animals, respectively (*n =* 51 syllables, *n =* 85 animals). Syllables are sorted by overall frequency within each syllable family. Animals are grouped by colony and ordered within each colony by similarity in syllable expression. (**E**) Transition graph of syllable families across all recordings. Node size indicates syllable-family frequency, and arrow width indicates transition frequency between and within syllable families. Transitions with frequencies below 5% are not shown. See also fig. S1B for individual colony visualizations.

We generated a UMAP embedding of observed syllable instances using latent pose-trajectory features inferred by Keypoint-MoSeq, revealing graded variation within and among syllable families (Fig. 1B; Supplementary Video 2). This variation reflected differences in kinematic features, including forward displacement, head turning, speed, and duration (Fig. 1C). For instance, forward-movement syllables were distinguished from other syllable families primarily by average speed and duration, with a smaller contribution from head turning. In contrast, scratch syllables had kinematic properties similar to those of turn syllables, but the latent MoSeq features captured subtle differences between these families in the embedding space. Animals showed variation in their relative use and sequencing of syllables (Fig. 1D and E; fig. S1B). In the subsequent sections, we leveraged this variation to ask whether it predicted key features of naked mole-rat societies.

### Spontaneous behavior predicts colony membership and individual identity

Naked mole-rat colonies are cohesive social units with colony-specific vocal and odor signatures (19, 21). We asked whether such colony-level variation is also reflected in the fine-scale behavioral syllables expressed outside social interactions. We trained a random forest classifier on behavioral syllable profiles from all but one animal and predicted the colony of the held-out animal (Fig. 2A). The classifier correctly predicted colony membership for 56 of 85 animals (65.9%; *P* = 5.0 × 10^−6^ by label permutations; Fig. 2B). Prediction exceeded chance in every colony (63.2–66.7% versus 25%; all adjusted *P* < 0.01, one-sided exact binomial tests with Bonferroni correction). Thus, spontaneous behavior recorded in isolation carries colony-specific signatures.

**Figure 2.**
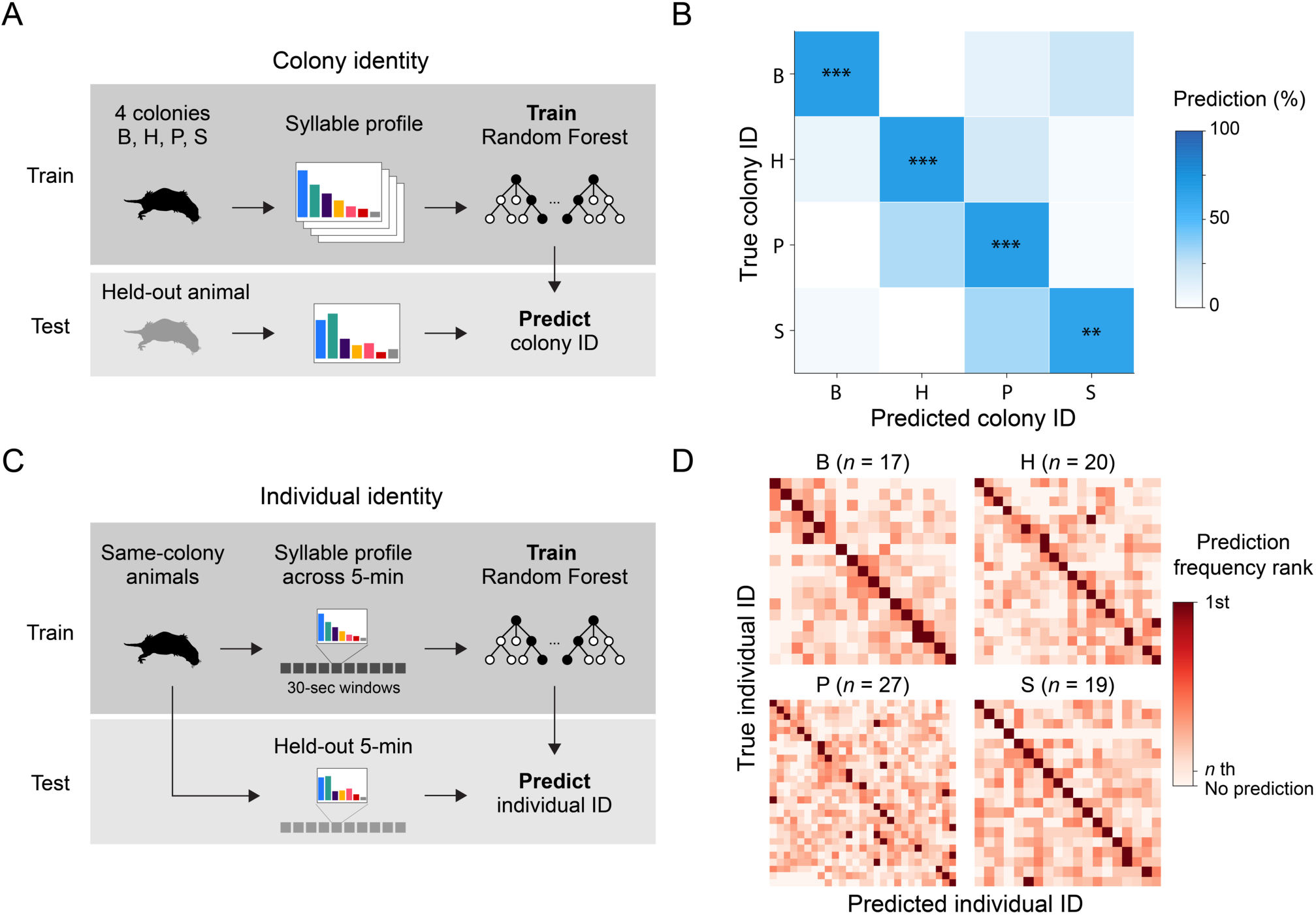
Spontaneous behavior predicts colony and individual identity. (**A**) Schematic of colony identity prediction. (**B**) Colony identity prediction results. Color indicates the percentage of predictions assigned to each colony. Asterisks indicate prediction rates above chance (** and ***, Bonferroni-adjusted *P* < 0.01 or < 0.001, respectively, by one-sided exact binomial tests). (**C**) Schematic of individual identity prediction. (**D**) Individual identity prediction results. Color indicates prediction-frequency rank within each true-animal row. Within each colony, animals are ordered by similarity in syllable expression. Identity prediction exceeded chance in every colony (Bonferroni-adjusted *P* < 0.001, one-sided exact binomial tests across four colonies).

Individual naked mole-rats also show stable, task-related behavioral differences, varying in their contributions to pup care, work, and colony defense (20, 23; but see 24, 25). We therefore reasoned that spontaneous behaviors may also carry individual-specific signatures. To test this hypothesis, we analyzed each colony separately and divided every animal’s 10-minute recording into twenty 30-second windows (Fig. 2C). We held out the same consecutive 10-window block from every animal, trained a random forest classifier on the remaining 10 windows from all animals in that colony, and predicted animal identity for each held-out window. We repeated this analysis across all possible positions of the held-out block. Syllable profiles strongly predicted individual identity in three colonies of 17–20 animals (Fig. 2D). The true identity received more predictions than any other identity for 88.2% of animals in B (15 of 17, adjusted *P* = 1.70 × 10^−16^; one-sided exact binomial tests against 1/*N* chance, with Bonferroni correction), 80.0% in H (16 of 20, adjusted *P* = 2.44 × 10^−17^), and 89.5% in S (17 of 19, adjusted *P* = 1.13 × 10^−19^). Performance was more modest but remained well above chance in the larger colony P (55.6%, 15 of 27, adjusted *P* = 1.54 × 10^−14^). The results were robust to both the analysis window size and the train–test proportion (fig. S2).

Reproductive status is a major axis of differentiation in naked mole-rat colonies, whereas nonbreeding males and females show little sexual dimorphism (26). We asked whether spontaneous syllables could also predict sex or reproductive role. Sex prediction did not exceed chance in any colony (fig. S3; balanced accuracy per colony = 37.7–63.9%, all Holm-adjusted *P* > 0.05; one-sided within-colony label-permutation tests). Similarly, reproductive-status predictions did not exceed chance. A three-class model distinguishing queens, breeding males, and nonbreeders and a binary model distinguishing breeders from nonbreeders both performed no better than chance (both Holm-adjusted *P* > 0.05; one-sided within-colony label-permutation tests). Together, these analyses show that spontaneous behavioral syllables carry specific rather than generic social signatures: they do not reflect sex or reproductive status, yet robustly signal both colony and individual identity.

### Naked mole-rat social hierarchy is linear and stable

Dominance hierarchies are a widespread organizing feature of animal societies. By structuring access to resources and reproductive opportunities, hierarchies can limit costly conflict and stabilize groups (2, 27). Naked mole-rat societies provide a prominent example in which dominance shapes both routine agonistic interactions and reproductive succession. Breeders initiate most shoves, and shoving among nonbreeders also covaries with dominance rank (28–30). When a queen dies or is removed, high-ranking nonbreeding females often compete intensely for the vacant breeding position (28), although peaceful succession has also recently been documented (31). We therefore first established the rank order within each colony and asked whether spontaneous behavioral syllables predicted those ranks.

To establish colony-wide rank orders for subsequent behavioral prediction, we used a custom tube test based on crawl-over interactions, similar to those used in previous studies (19, 32; Methods). Two animals were introduced to meet head-on in a narrow tube, and the animal that crawled over its opponent was scored as the winner (Fig. 3A). Crawl-over behavior was fast and stereotyped: following the initial face-to-face touch, the subordinate animal adopted a posture that allowed the dominant animal to crawl over it, and the entire sequence typically took under a second (Supplementary Video 3). We repeated this test across dyads and trials to derive each animal’s rank within its colony. Rank derived from crawl-over behavior has been shown to track dominance inferred from agonistic behaviors such as shoving (28, 29).

**Figure 3.**
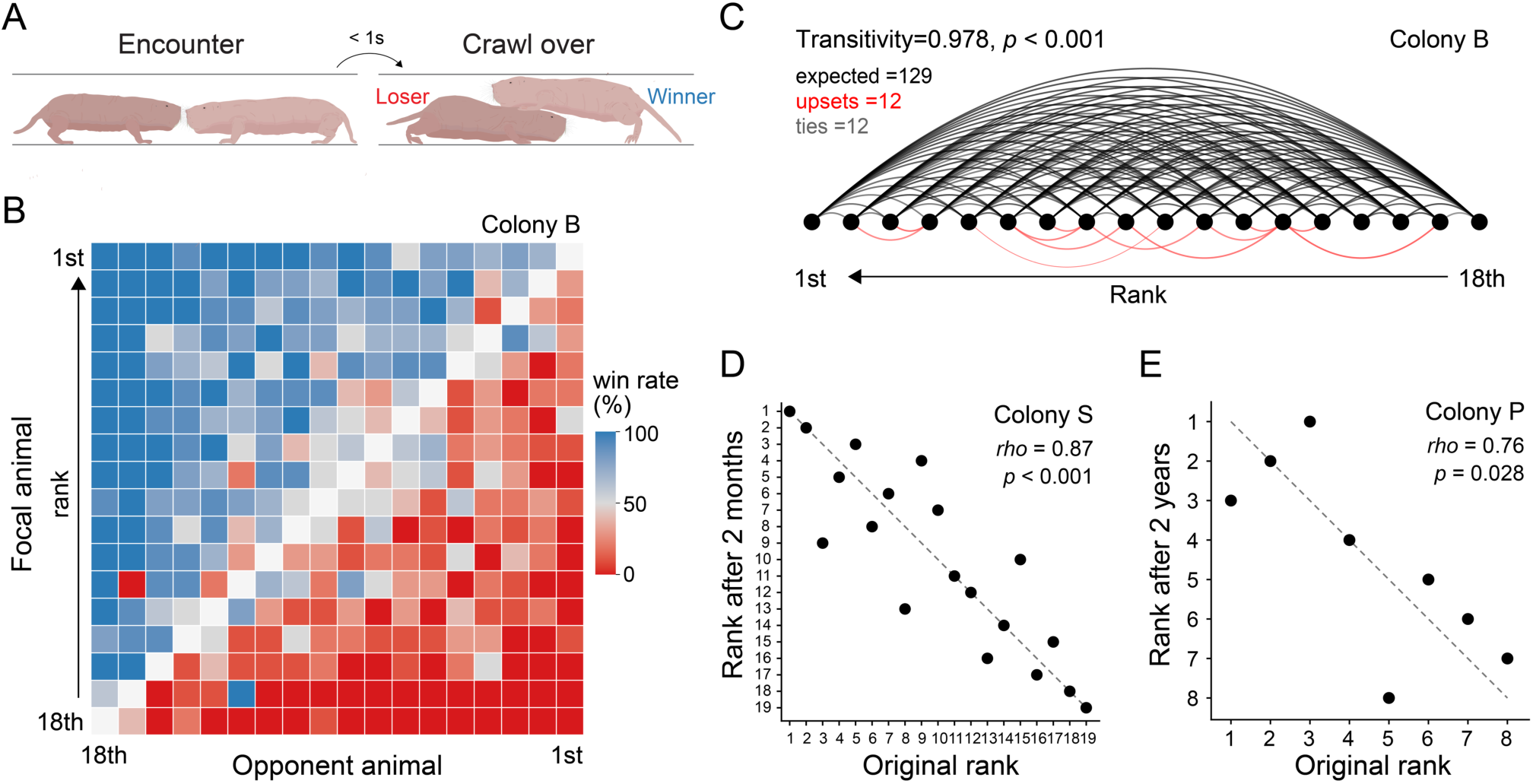
Naked mole-rat social hierarchy is linear and stable. (**A**) Schematic of the tube test used to determine dominance relationships between naked mole-rats. See also Supplementary Video 3. (**B**) Matrix of tube-test results in colony B. Each cell shows the focal animal’s win rate across 10 repeated tube-test trials against an opponent. (**C**) The hierarchy is highly transitive. Circles represent animals in colony B, and each line represents the tube-test outcome for a dyad. Black lines (*n* = 129) indicate outcomes consistent with the Bradley–Terry rank order, in which the higher-ranked animal defeated the lower-ranked animal, whereas red lines (*n* = 12) indicate upsets. Ties (*n* = 12) are not shown. Transitivity is the proportion of fully tested, untied triads with noncyclic dominance relationships. The *P* value was calculated using a permutation test that randomized trial outcomes within dyads under equal win probabilities. See fig. S5 for other colonies. (D and E) Ranks remained stable over 2 months in colony S (**D**) and over 2 years in colony P (**E**). *rho* and *P,* Spearman’s rank correlation test.

Because exhaustive, repeated testing of every possible dyad becomes prohibitive as colony size increases, we used a Bradley–Terry model to infer latent dominance scores and colony-wide ranks from a connected subset of dyads (Fig. 3B). We evaluated the accuracy and robustness of this approach in simulations that subsampled dyads and trials from a fully sampled dataset from colony B. These simulations identified sampling regimes that recovered the full-data rank structure without exhaustive all-by-all testing (fig. S4; Methods).

We ran the tube test to rank the animals from the four colonies and found that naked mole-rat dominance hierarchies were highly linear, consistent with previous findings (28). In colony B, where all possible dyads were tested, only 12 of 141 untied dyads were upsets relative to the rank order. Transitivity was significantly higher than expected by chance (Fig. 3C; fraction of triads with transitive relationships = 0.978, *P* < 0.001 by permutation test). The other three colonies whose dyads were partially sampled showed similarly high transitivity, with 94.7–97.9% of triads being noncyclic (all *P* < 0.001; permutation tests; fig. S5).

Although rank has been routinely characterized in naked mole-rat colonies (19, 28, 32), how long colony-wide rank order persists in naked mole-rats remains unclear. We repeated rank measurements at several intervals. Remarkably, we found that ranks remained highly consistent after 2 months in colony S (Spearman’s *rho* = 0.87, *P* < 0.001, *n* = 19; Fig. 3D), after 4 months in colony H (*rho* = 0.77, *P* < 0.001, *n* = 18; fig. S6), and even after 2 years among the eight animals characterized at both time points in colony P (*rho* = 0.76, *P* = 0.028; Fig. 3E). Together, these results show that naked mole-rat colonies form highly linear hierarchies whose rank order can persist for months to years.

### Spontaneous behavior predicts social rank

Having established stable colony rank orders, we next asked whether spontaneous behavior in isolation could predict an animal’s social rank. We used partial least squares (PLS) regression (33), which summarizes behavioral profiles into a few latent components that covary with rank (see Methods for details). Within each colony, we held out each pair of animals in turn and regressed the relative ranks of the remaining animals on their syllable profiles (Fig. 4A, left). The fitted model then scored both held-out animals, and the animal with the higher score was predicted to be dominant (Fig. 4A, right). Neither member of a test dyad contributed to model fitting. These predicted dyadic dominance relationships were then used to reconstruct each colony’s rank order.

**Figure 4.**
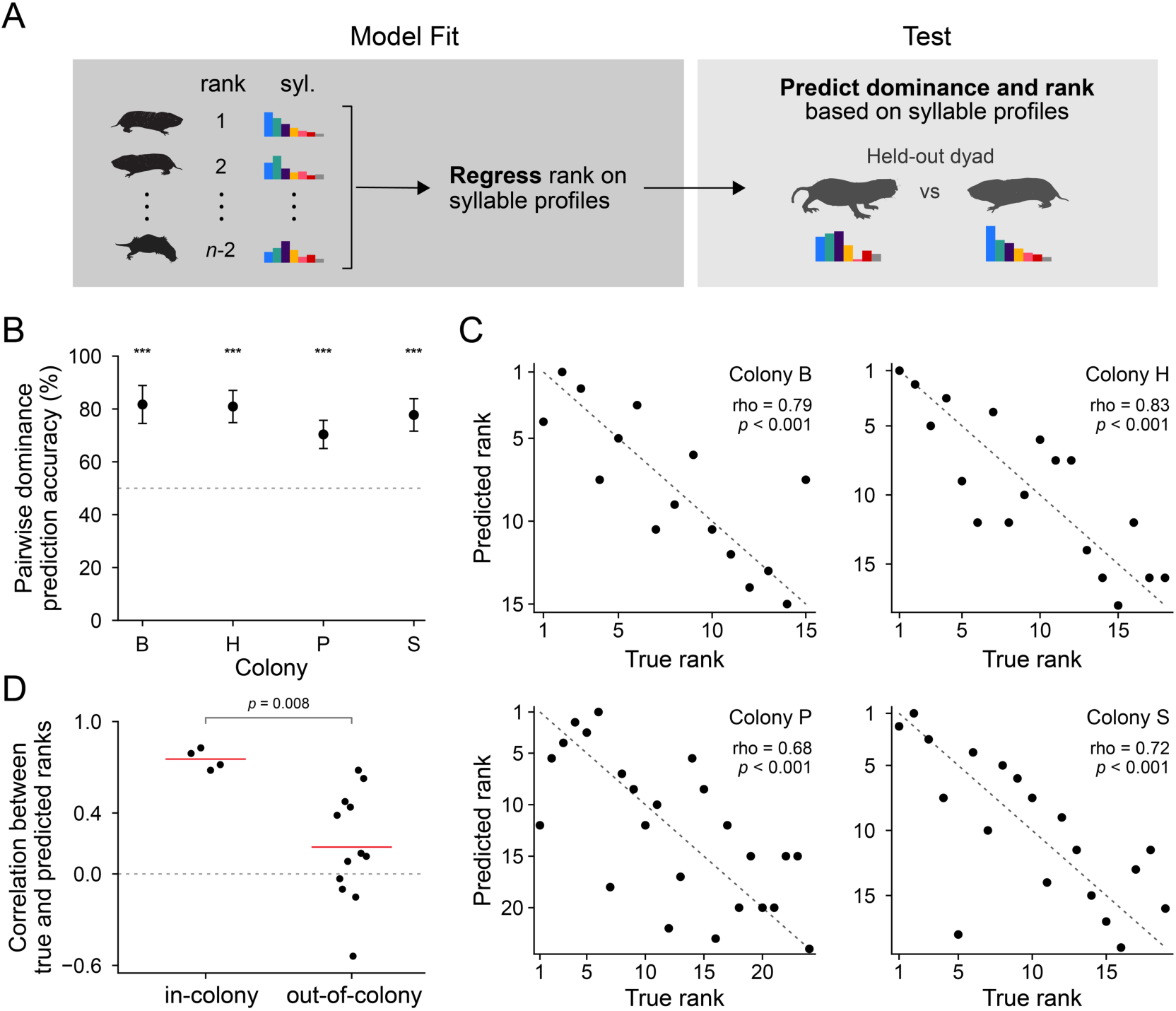
Spontaneous behavior predicts social rank. (**A**) Schematic of rank prediction. Within each colony, each pair of animals was held out in turn, and a partial least squares (PLS) regression of relative rank on syllable profiles was fitted to the remaining animals. The fitted model scored both held-out animals, and the animal with the higher score was predicted to rank higher. Predictions across all held-out pairs were combined to reconstruct the within-colony rank order. (**B**) Pairwise rank prediction accuracy across colonies. Error bars indicate Wilson 95% binomial confidence intervals, and the dashed line indicates chance accuracy (50%). ***, Bonferroni-adjusted *P* < 0.001 by one-sided exact binomial tests against chance. (**C**) Predicted versus true ranks across colonies. *rho and P,* Spearman’s rank correlation test. (**D**) Syllable-based rank prediction is colony-specific. Correlations between true and predicted ranks are shown for in-colony predictions (*n =* 4), where training and test animals came from the same colony, and out-of-colony predictions (*n =* 12), where they came from different colonies. Each point represents one training–test colony combination; red lines indicate means. *P*, two-sided exact source-stratified permutation test. See also fig. S7.

Strikingly, pairwise dominance was predicted well above chance in every colony (Fig. 4B), with accuracies of 82.9% in colony B, 81.7% in H, 70.7% in P, and 78.4% in S (all Bonferroni-adjusted *P* < 0.001, one-sided exact binomial tests). The pairwise predictions accurately recovered colony rank orders (Fig. 4C): predicted and true ranks were strongly correlated across colonies (Spearman’s *rho* = 0.79 in colony B, 0.83 in H, 0.68 in P, 0.72 in S; *P* < 0.001 for all colonies). Finally, we asked whether the predictive power of behavioral syllables for rank generalized across colonies. Rank prediction was substantially stronger when the training and test animals came from the same colony than when they came from different colonies (mean *rho* = 0.75 versus 0.18; *P* = 0.008, two-sided exact source-stratified permutation test; Fig. 4D and fig. S7). Two of the 12 cross-colony transfers were nonetheless successful: models fitted in colony P predicted rank in S, and models fitted in S predicted rank in H, suggesting rank–behavior patterns shared among some but not all colonies (fig. S7). Together, these results show that social rank is expressed through behavioral patterns that are largely specific to each colony.

## DISCUSSION

Animal societies persist over months to years, yet they are built from individual actions lasting fractions of a second. Our results show that in naked mole-rats, these scales are tightly linked. Fine-scale behavioral syllables in solo recordings predicted individual identity and colony membership. They also predicted positions within dominance hierarchies that are stable over months to years. Spontaneous behavioral syllables in naked mole-rats therefore reflect multiple scales of their enduring social organization.

Previous studies across vertebrates and social insects have linked specific individual behavioral tendencies to roles within groups and to collective performance (12–15). In house finches, exploration in a novel environment predicted the number of social partners in the wild (12). In sticklebacks, individually measured sociability and exploration predicted spatial position, leadership, movement dynamics, and foraging performance within shoals (13). In ants, exploration by isolated workers was associated with foraging roles and the performance of experimentally assembled groups (14, 15). Our high-resolution behavioral study extends this cross-scale link to a mammalian society. Beyond naked mole-rats, studying other African mole-rats would provide a natural comparative system as closely related species range from solitary to eusocial systems (16, 34, 35). Applying similar approaches across related mole-rat species would reveal how the information carried by individual spontaneous behavior changes with social complexity.

Our finding that behavioral syllables carried individual and colony signatures (Fig. 2) parallels recent studies of social communication via vocalizations and odor (19, 21). Soft chirps, a frequent naked mole-rat vocalization, encode both individual and colony signatures (19), and odor profiles likewise distinguish colonies (21). Theory predicts that overlapping information across multiple sensory modalities can make communication more robust to noisy or changing conditions (36, 37). Such redundancy of social information occurs across taxa (37–40). Thus, individual identity and colony membership are reflected in movement, sound, and odor, potentially making social recognition robust when only one channel is reliable.

Alongside this redundancy, each modality also appears to carry information that the others do not. Vocalizations form a large repertoire with context-specific calls for greeting, courtship, alarm, and aggression (16, 17), and colony dialects are learned and stabilized by the queen, making sound a vehicle for colony cohesion and vocal culture (19). Odor is tied most closely to reproduction: a queen-specific ester suppresses reproduction in nonbreeders and governs queen succession (21). Behavioral syllables, by contrast, did not predict sex or reproductive status (fig. S3). Instead, our results show that behavioral syllables carried predictive information about dominance rank (Fig. 4). Rank has been comparatively difficult to read out from other channels: soft-chirp features did not strongly predict rank (19), and odor profiles distinguish colonies and queens but have not been linked to dominance rank (21). Behavioral syllables may thus be a channel in which dominance rank is particularly legible, complementing the identity, cultural, and reproductive information conveyed by sound and odor.

How colony mates access rank information carried by movement remains an open question. Naked mole-rats are functionally blind and interact through frequent physical contact, making somatosensation a plausible channel (17). Deflecting a single body hair triggers precise orientation, and nearly one-third of primary somatosensory cortex represents the incisors used in social interactions, suggesting naked mole-rats have a heightened ability to process somatosensory information (41, 42). Relatedly, indirect communication via substrate vibration may provide another route. The solitary Cape mole-rat (*Georychus capensis*), another bathyergid, transmits mate-attraction signals between neighboring burrows by alternately drumming its hind feet on the burrow floor (43). Naked mole-rats could likewise transmit rank-related information through substrate-borne vibrations. Altogether, we speculate that naked mole-rats may detect rank-predictive locomotion patterns of conspecifics through direct or indirect somatosensory communication. Future studies could test this hypothesis, for example, by controlled encounters with tactile and vibratory perturbations.

One limitation of our study is that body weight and rank are closely associated in naked mole-rats (28), so their contributions to behavioral prediction could not be fully separated. When we restricted the analysis to pairs of animals differing in body weight by no more than 10%, behavioral syllables still predicted the dominant animal in 75–79% of cases in colonies B, H, and S, although accuracy fell to 59% in colony P. Thus, behavior carries rank information among animals of similar weight, although weight-related variation may also contribute to prediction. Future studies that sample additional colonies, particularly colonies with narrower body-weight ranges, could provide more comparisons between similarly weighted animals of different ranks and further separate these effects.

Finally, the behavioral signature of rank in naked mole-rats provides a tractable readout for studying how the brain represents complex social hierarchy. Studies in mice and other animals have begun to identify neural mechanisms that encode and regulate dominance (27, 44). Yet most experiments have examined pairs or small groups largely because of biological constraints in model organisms. Naked mole-rats are poised to extend this line of inquiry to a large mammalian society with a stable hierarchy. Modern neuroscience methods, such as functional ultrasound imaging and Neuropixels, have increasingly been applied to mole-rats (21, 45). Combining these tools with quantitative tracking across solo and social contexts provides an exciting avenue for understanding how neural activity generates rank-associated behavior and dominance hierarchies in complex vertebrate societies.

## MATERIALS AND METHODS

All procedures were performed in accordance with the National Institutes of Health Guide for the Care and Use of Laboratory Animals and were approved by the Columbia University Institutional Animal Care and Use Committee, protocol number AC-AABP9550.

### Animals

Four captive colonies, named B, H, P, and S, were used for this study. Animals were maintained in a climate-controlled room at 80–85°F and approximately 50% relative humidity on a 12-hour light/12-hour dark cycle. Colonies contained animals of both sexes and were housed in interconnected mouse- and rat-cage habitats joined by clear acrylic tubing. The number and arrangement of chambers were adjusted to colony size. Animals received sweet potato, celery, and apple daily, supplemented twice weekly with vitamin-enriched cereal. Animals were identified using implanted radio-frequency identification transponders and were weighed regularly.

### Recording

Animals were recorded individually for 10 minutes in a circular polyethylene arena with a 43.2 cm inner diameter (Fig. 1A; United States Plastic Corp., catalog no. 14317). The floor was covered with an approximately 6-mm layer of black terrarium sand (ReptiSand Midnight Black, Zoo Med). A monochrome Blackfly S USB3 camera (BFS-U3-13Y3M-C, FLIR) fitted with an 8-mm fixed-focal-length lens (33-307, Edmund Optics) was mounted above the center of the arena. Videos were acquired in SpinView at 60 frames per second and 1,280 × 1,024-pixel resolution. At the start of each recording, the animal was placed in the center of the arena and allowed to move freely without a conspecific present. Recordings were performed in the room in which the animals were housed. Before recording, each animal was handled for at least 3 days and habituated to the arena for at least 10 minutes on each of 3 days. Recordings from different colonies were interleaved across days and spread across the working day. All recordings used the same arena, camera and acquisition settings. Sand was reused within a colony and replaced between colonies, and the arena was cleaned with ethanol and water between sessions. The analyzed dataset contained 85 recordings, one recording per animal (B, *n =* 18; H, *n =* 21; P, *n =* 27; S, *n =* 19), comprising 3.45 million frames (Fig. 1A; fig. S1A).

### Pose inference

We performed pose inference across frames with SLEAP (4) using a seven-landmark skeleton: nose, left and right ears, body centroid, left and right hind paws, and tail base (Fig. 1A). Approximately 5,000 frames were manually labeled to train a top-down pose-estimation model. We then used the trained model to predict body-part locations in all frames. We treated keypoints with prediction confidence below 0.5 or identified as spatial outliers using an outlier scale factor of 6.0 as missing. We linearly interpolated their coordinates across neighboring frames. We visually inspected the resulting pose trajectories.

### Behavioral syllable inference

We inferred behavioral syllables from the pose trajectories with Keypoint-MoSeq (8). We used principal component analysis to summarize coordinated movements of the seven keypoints with a smaller set of pose variables. The first five principal components explained 92.1% of the pose variance, so we represented pose dynamics using five latent dimensions. We used three autoregressive lags to identify recurring movements over time. We allowed the model to infer up to 100 candidate behavioral syllables. To select the temporal scale kappa, we fit models across kappa values spanning 10^5^ to 10^7^ and compared syllable-duration distributions, syllable usage, representative trajectories and videos. We found that models with kappa of 10^7^ consistently produced syllables without excessive fragmentation and therefore used it for the final model. At kappa = 10^7^, we then fit 30 randomly initialized models. Each model underwent 50 iterations of autoregressive hidden Markov model initialization followed by full model fitting through iteration 500. We selected the model with the highest estimated marginal likelihood as the final model.

The final Keypoint-MoSeq model contained 67 behavioral syllables with at least 25 instances across videos. We took these syllables as an initial candidate set of naked mole-rat spontaneous behavioral syllables. We generated representative video grids and keypoint-trajectory summaries for the candidate syllables and curated them based on movement type, direction, speed, and confidence statistics. After careful inspection, 16 candidate syllables were excluded because their instances were inconsistent: each bundled qualitatively different movements under a single syllable label; in preliminary UMAP embeddings, instances of these syllables were also scattered across behavioral space rather than forming distinct clusters, together suggesting that they represented noise rather than coherent but unnamed behaviors. The remaining 51 behavioral syllables with consistent and interpretable movement patterns expressed across videos were retained as high-confidence syllables and grouped into seven broad families: forward movement, reverse movement, pause, scratch, dig, turn, and stand up (Fig. 1A; fig. S1A). Supplementary Video 1 shows example movies for the syllables.

We defined a syllable instance as a contiguous run of frames assigned to the same behavioral syllable. The final curated dataset contained 50,728 syllable instances across 3,234,486 frames, corresponding to 93.6% of all frames. The fraction of frames assigned to high-confidence syllables was similar across recordings (mean ± SD, 93.8% ± 4.8%) and across colonies (B, 93.1%; H, 92.0%; P, 94.2%; S, 96.0%). For each animal, we calculated the usage of each of the 51 high-confidence syllables as the fraction of high-confidence frames assigned to that syllable (Fig. 1D). Five additional summary statistics were calculated for each animal: mean absolute angular velocity, number of syllables observed, mean and maximum syllable-instance duration, and number of unique syllable transitions. All classification analyses below used the 51 syllable fractions and five summary statistics.

### Instance-level UMAP embedding

To map the behavioral space of behavioral syllables expressed by naked mole-rats recorded alone, we performed dimensionality reduction using uniform manifold approximation and projection (UMAP) (Fig. 1B and C). Each dot in the UMAP represents one syllable instance lasting at least 10 frames (167 ms). The input features were the five-dimensional latent pose trajectory inferred by Keypoint-MoSeq, forward displacement, lateral displacement, and heading change. We resampled each trajectory to 25 normalized time bins and standardized all features. We then fit the UMAP using Euclidean distance, 30 neighbors, and a minimum distance of 0.2. An example movie for behavior-to-UMAP correspondence is available (Supplementary Video 2).

### Colony-membership prediction

We predicted colony membership from the syllable profile of each animal (*n =* 85) using leave-one-animal-out cross-validation. In each fold, a random forest classifier with 50 trees, maximum depth 8, and balanced-subsample class weights was trained on the other 84 animals and used to classify the held-out animal. We assessed overall significance by permuting colony labels 200,000 times and compared the observed leave-one-animal-out accuracy with the resulting null distribution. Prediction accuracy was tested against 0.25 chance with one-sided exact binomial tests and Bonferroni correction across the four colonies.

### Individual-identity prediction

To predict individual identity, we analyzed each colony separately. We divided each animal’s 10-minute recording into 20 nonoverlapping 30-second windows. Each window contained 1,800 frames and was represented by its 56-feature behavioral syllable profile. For the primary analysis, we used a continuous 5-minute block as the test set and the remaining 5 minutes as the training set. We shifted the 5-minute test block forward in 30-second steps, producing 11 overlapping test blocks and 11 train–test splits per colony. For each split, we trained a random forest to associate the behavioral profiles of the training windows with animal identity. The classifier used 20 trees, a maximum depth of 6, and balanced-subsample class weights. We then predicted identity from the behavioral profile of each window in the test block. To determine whether an animal’s identity was recovered, we pooled its predictions across all 11 splits. This produced 110 window-level predictions per animal. We considered identity recovered when the true identity was the single most frequently predicted label. We compared the number of animals recovered with a chance probability of 1/*N*, where *N* was the number of animals in the colony. We used one-sided exact binomial tests with Bonferroni correction across colonies.

To determine whether identity prediction depended on analysis choices, we conducted two sensitivity analyses (fig. S2). First, we tested the effect of window size. We divided each recording into nonoverlapping 15-, 30-, 45-, or 60-second windows and calculated a 56-feature behavioral profile for each window. We retrained the classifier for every window size and split. We used 5-minute training and 5-minute test sets and moved the 5-minute block across the recording in one-window steps. We excluded windows that crossed a training–test boundary. Second, we tested the effect of training and test duration. Using 30-second windows, we evaluated training durations of 5:5, 6:4, 7:3, 8:2, 9:1, and 9.5:0.5 minutes. For each ratio, we moved a contiguous test block across the recording and used all remaining windows for training. These sensitivity analyses found that individual identity prediction was robust to analysis window size and training–test ratio (fig. S2).

### Sex and reproductive status prediction

To predict sex, we included 82 animals with known sex (B, *n =* 18; H, *n =* 21; P, *n =* 26; S, *n =* 17). We trained separate classifiers within each colony. We also pooled the held-out predictions from all four colonies to obtain a combined performance estimate (fig. S3). To test whether reproductive status influenced sex prediction, we repeated the per-colony analysis after excluding queens and breeding males. None of the per-colony sex predictions was significant after correction, either with or without breeders. To predict reproductive status, we analyzed all 85 animals across colonies because each colony contained only one or two breeders. We evaluated a three-class model that distinguished queens, breeding males, and nonbreeders. We also evaluated a binary model that distinguished breeders from nonbreeders (fig. S3). Neither model performed above chance.

For both sex and reproductive-status predictions we used the 56-feature behavioral profile of each animal and the leave-one-animal-out procedure for all classifiers. In each fold, we trained a random forest on all but one animal and predicted the held-out animal. Each classifier used 50 trees, a maximum depth of 8, and balanced-subsample class weights. To assess significance, we shuffled labels within each colony 1,000 times and repeated the complete leave-one-animal-out analysis. We calculated one-sided permutation *P* values from balanced accuracy with Holm correction.

### Tube-test assay and rank inference

#### Tube-test assay

We measured pairwise dominance using a tube-test assay informed by previous tube-test protocols used in the naked mole-rat (19, 32). We removed two animals from their colony and gently placed them at opposite ends of an acrylic tube identical to those used in their laboratory habitat. The animals then approached each other head-on. Shortly after their initial face-to-face contact, one animal adopted a yielding posture that allowed the other to crawl over it. This stereotyped interaction occurred rapidly, typically within 1 second (Supplementary Video 3). We scored the animal that crawled over its opponent as the winner of the trial and tested each dyad for 3–10 trials (see *Exploratory analyses for an effective sampling scheme* for how we determined the number of trials per dyad).

#### Bradley–Terry rank estimation

We estimated dominance relationships from tube-test results by a ridge-regularized Bradley–Terry model (46). The model uses pairwise win–loss outcomes to place animals on a common dominance scale. A strength of the method is that it can estimate ranks from a subset of dyads, provided that the dyad network is connected. This property allowed us to efficiently estimate colony-wide rank.

For animals *i* and *j* with latent dominance scores *θ_i_* and *θ_j_*, the probability that *i* defeated *j* was

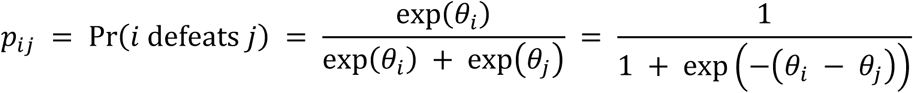

Let *w_ij_* and *w_ji_* denote the observed numbers of wins in each direction for a tested dyad. We estimated the scores by maximizing the ridge-penalized log likelihood

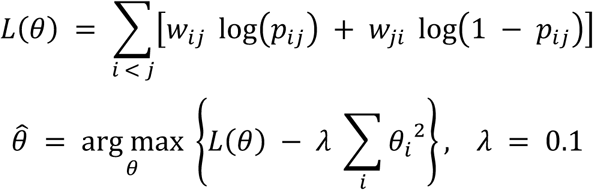

During model fitting, we selected one animal as an arbitrary reference and fixed its latent dominance score at zero. After fitting, we subtracted the colony mean from each animal’s latent dominance score so that the mean score within each colony was zero. We then ranked animals from highest to lowest dominance score.

#### Exploratory analyses for an effective sampling scheme

We next asked how many dyads and trials per dyad were needed to recover colony-wide rank reliably. A colony of *n* animals contains 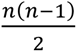 possible dyads. Thus, colonies of 20–30 animals contain 190–435 possible dyads. Ten trials for every dyad would require 1,900–4,350 tube-test trials for one complete census. Repeating this design across several colonies and time points would become prohibitive as colony sizes increase. We therefore collected an exhaustive reference dataset and used retrospective simulations to choose the effective number of dyads and trials per dyad before testing the remaining colonies.

To establish the full-data reference, we exhaustively tested colony B. This colony contained 18 animals and 153 possible dyads. We tested every dyad for 10 consecutive trials, which produced 1,530 trial outcomes. We fitted the Bradley–Terry model to all outcomes to define the full-data rank order. We then retrospectively subsampled this dataset. For every simulated condition, we refitted the Bradley–Terry model and compared the subsampled rank order with the full-data reference using Spearman’s rank correlation. We ran 1,000 simulations per condition (fig. S4).

To evaluate sampling effort, we designed an adaptive procedure informed by body weight. Body weight is readily available before testing and is correlated with rank in naked mole-rats (28, 29). We therefore began each simulated network with a round-robin among weight-stratified anchor animals. We placed the remaining non-reproductive animals by weight-guided bracketing against animals near their provisional ranks. We tested reproductive animals separately against high- and mid-ranked comparators because their ranks tend not to correlate as clearly with weight as nonbreeders (47). We then added local refinement dyads between neighboring provisional ranks. We kept the network connected throughout. Body weight guided dyad selection but was not included in the Bradley–Terry model.

We used two complementary analyses to evaluate the number of trials per dyad and the proportion of dyads tested (fig. S4). We considered the results together when selecting the final sampling scheme. First, to evaluate the number of trials per dyad, we varied the number of trials from 1 to 10 while holding dyad coverage constant at 40% (fig. S4A). For each selected dyad, we sampled outcomes without replacement from its 10 observed outcomes. Mean rank correlation increased from 0.837 with one trial per dyad to 0.936 with three trials and 0.960 with five trials. Gains became much smaller after three trials. Based on these results, we devised a “3–0 or 5” rule: testing stopped after a 3–0 sweep; otherwise, testing continued to a maximum of five trials. This rule provided more evidence than fixed three-trial testing while requiring fewer trials than fixed five-trial testing. Second, to evaluate dyad coverage, we applied the “3–0 or 5” rule and varied the proportion of possible dyads tested from approximately 10% to 100% (fig. S4B). Mean rank correlation with the full-data reference reached 0.92 at approximately 30% dyad coverage, 0.95 at 40%, and 0.97 at 50%, with diminishing improvements at higher coverage.

Together, our analyses of colony B supported testing 30%–50% of possible dyads and 3–5 trials per dyad. This approach required approximately 12%–20% of the trials needed to test every dyad 10 times and still recovered the full-data rank order accurately (Spearman’s *rho* = 0.92–0.97). We therefore used this sampling scheme to rank colonies H, P, and S (Fig. 3; figs. S5 and S6).

### Hierarchy linearity and temporal stability

To quantify the linearity of the dominance hierarchy in each colony, we assigned dominance in each tested dyad to the animal that won more trials. We excluded tied dyads from this analysis. We defined a complete triad as three animals for which all three constituent dyads were tested and untied. We calculated transitivity as the fraction of complete triads that did not form a three-animal dominance cycle. To determine whether the observed transitivity exceeded chance, we performed 10,000 permutations. In each permutation, we held the observed number of trials for each dyad constant and redrew wins from a binomial distribution that assigned both animals an equal probability of winning. We then reassigned dyad directions and recalculated transitivity using the same criteria. We calculated the one-sided permutation *P* value as (*k* + 1)/10,001, where *k* was the number of randomized datasets with transitivity equal to or greater than the observed value. Thus, smaller *P* values indicated that the observed hierarchy was more transitive than expected by chance (Fig. 3C; fig. S5).

To assess temporal stability of rank, we analyzed colonies S, H, and P, for which we collected tube-test data at two time points. We calculated their relative ranks at each time point and compared the two rank orders using a two-sided Spearman’s rank correlation test. We compared 19 animals in colony S across 2 months (Fig. 3D), 18 animals in colony H across 4 months (fig. S6), and eight animals in colony P across 2 years (Fig. 3E). The colony P comparison included only eight animals because the first assay ranked a subset of 10 animals from the larger colony, and two of those animals died before the second, more comprehensive assay.

### Pairwise dominance and rank prediction

To predict relative dominance from spontaneous behavior, we fitted separate models within each colony (B, *n* = 15; H, *n* = 18; P, *n* = 24; S, *n* = 19). Each animal was described by the 51 syllable fractions and five summary statistics. To evaluate predictions for animals excluded from model fitting, we used leave-two-animals-out cross-validation, holding out each pair of animals in a colony in turn. In each fold, we standardized each feature using the mean and standard deviation of the remaining animals, re-ranked the remaining animals by their observed ranks, and scaled these relative ranks linearly from 0.5 (highest) to −0.5 (lowest). We then fitted a PLS regression of the scaled ranks on the standardized features (scikit-learn PLSRegression). We chose the number of latent components in PLS regression for each colony from leave-two-animals-out cross-validated rank prediction. The fitted model assigned a score to each held-out animal, and the animal with the higher score was predicted to rank higher (Fig. 4A). We pooled predictions across folds to calculate pairwise accuracy (Fig. 4B). We compared accuracy with 0.5 chance using one-sided exact binomial tests with Bonferroni correction across colonies. To reconstruct whole-colony rank (Fig. 4C), we counted, for each animal, the held-out pairs in which it was predicted to rank higher and ranked animals by this count. We compared the predicted and observed colony rank orders using two-sided Spearman’s rank correlation.

### Cross-colony transfer of rank prediction

To test whether the ability to predict rank from behavioral profiles within colonies generalized across colonies, we fitted the PLS regression model to all animals in one source colony, using the source colony’s feature standardization and number of components, and applied it to all animals in each of the other colonies. We then reconstructed each target colony’s predicted rank order from pairwise comparisons of predicted scores and compared it with the observed rank order using Spearman’s rank correlation. Together with the four within-colony results, this produced 16 source–target correlations. We compared the mean of the four within-colony correlations with the mean of the 12 cross-colony correlations using a two-sided exact permutation test. For each source colony, we treated each of its four target results in turn as a hypothetical within-colony result and treated the other three as cross-colony results. Selecting one result independently for each of the four source colonies produced 4^4^ = 256 assignments. The *P* value was the proportion of these assignments with an absolute within-versus-cross-colony difference at least as large as the observed difference (Fig. 4D; fig. S7). To evaluate individual transfers, we compared each of the 12 cross-colony correlations with a null distribution generated by permuting the observed ranks among target-colony animals 10,000 times, calculated one-sided *P* values, and adjusted them across the 12 source– target directions using the Holm method.

## ACKNOWLEDGMENTS

We thank members of the Abdus-Saboor lab for discussion. We thank Thomas Park, Rochelle Buffenstein, and Dan McCloskey for donating naked mole-rats to us and helping us to establish our own colonies. We thank Ewan St. John Smith for helpful suggestions on naked mole-rat husbandry. We also acknowledge support for building naked mole-rat habitats, behavioral setups, and housing from Tanya Tabachnik, Barbara Han, Marlon Lowe, and the Zuckerman Institute Advanced Instrumentation Core.

## FUNDING

This work was supported by the Leon Levy Foundation and the New York Academy of Sciences (Leon Levy Scholarship in Neuroscience to Y.H.) and the New York Hideyo Noguchi Memorial Society (NY Hideyo Noguchi Memorial Scholarship to Y.H.). I.A.-S. acknowledges support from the NIH Director’s Pioneer Award (DP1AT013723) and as a Freeman Hrabowski Scholar at the Howard Hughes Medical Institute.

## DATA AND CODE AVAILABILITY

Associated scripts and data are available on GitHub (github.com/YukiHaba/NMR_spontaneous_behavior).

## CONFLICT OF INTEREST

None declared.

---

**Supplementary Figure S1.**
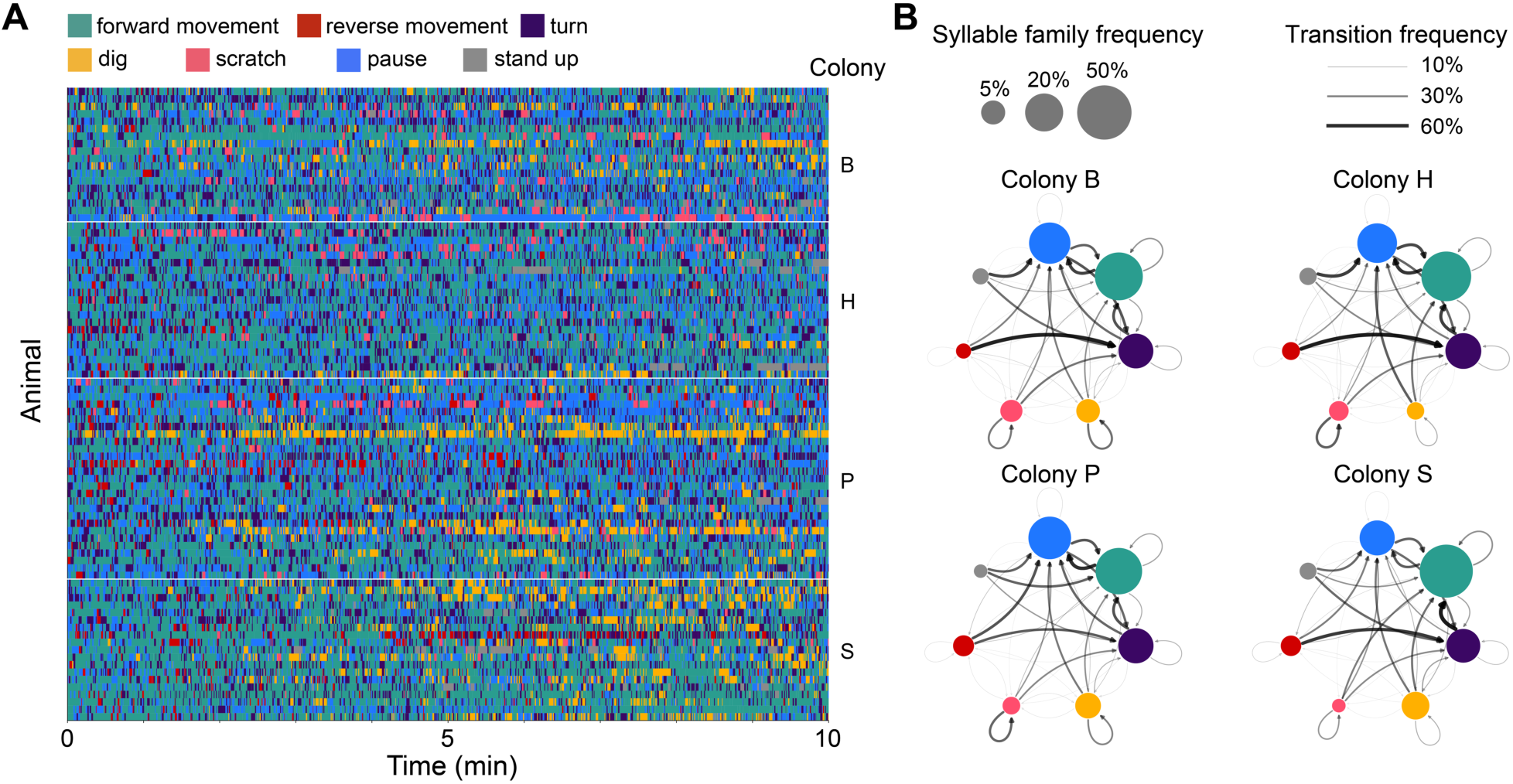
Additional details of spontaneous behavioral syllables, related to Figure 1. (**A**) Ethogram of expressed syllables in 10-minute recordings from 85 naked mole-rats in four colonies. Colors indicate syllable families. (**B**) Transition graphs of syllable families across recordings of the four colonies.

**Supplementary Figure S2.**
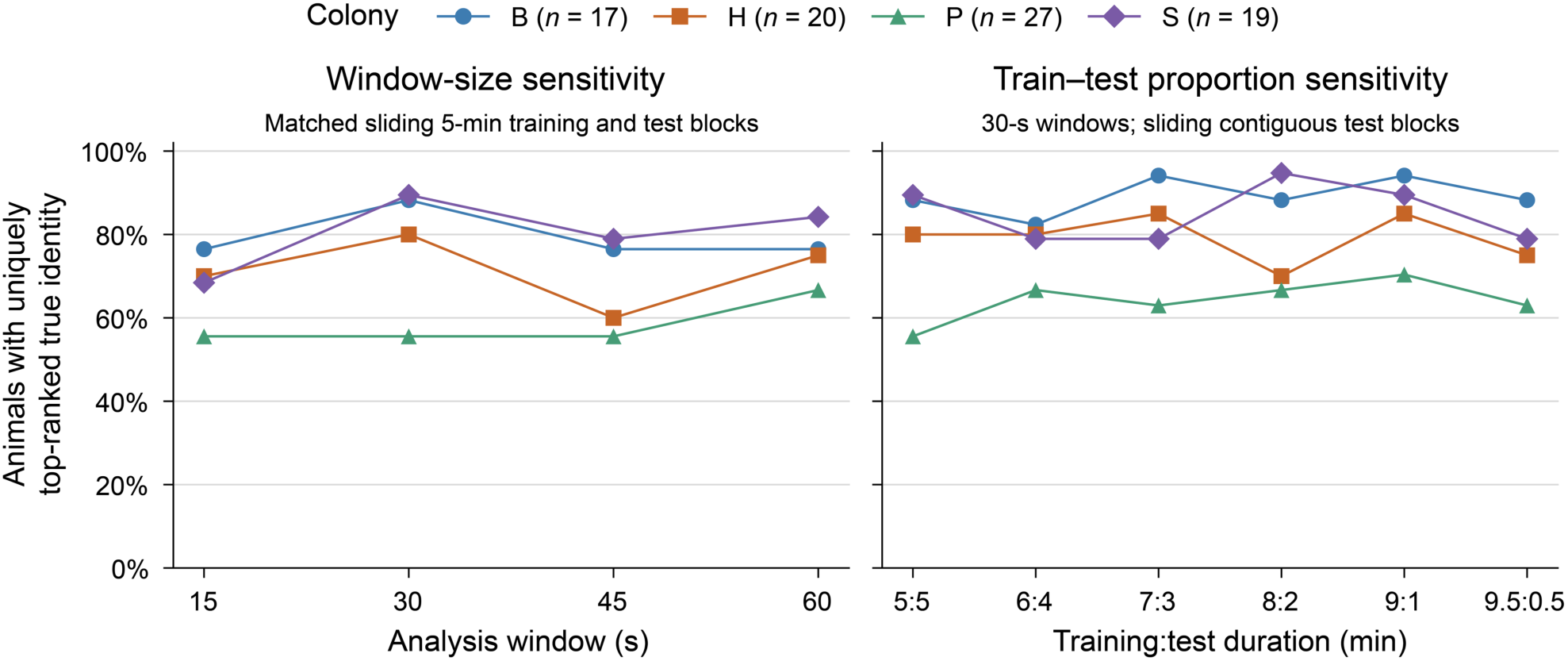
Individual identity prediction is robust to analysis window size and training–test ratio. (Left) Percentage of animals for which the true identity received more predictions than any other identity using analysis windows of 15, 30, 45, or 60 seconds. Classifiers were evaluated using matched sliding 5-minute training and test blocks. (Right) Percentage of animals with a uniquely top-ranked true identity across different training durations, using 30-second analysis windows and sliding contiguous test blocks. Colors and symbols indicate colony; sample sizes are shown in the key. Identity prediction remained above colony-specific (1/*N*) chance under every condition (all Bonferroni-adjusted *P* < 0.0001, one-sided exact binomial tests).

**Supplementary Figure S3.**
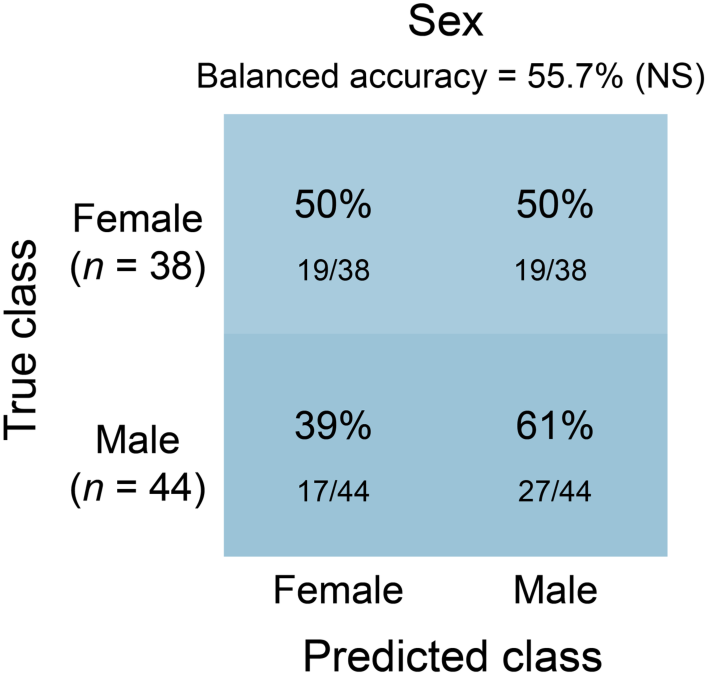
Spontaneous behavior did not predict sex. Row-normalized confusion matrix showing leave-one-animal-out sex predictions pooled across colonies. Random forest classifiers using syllable-frequency and behavioral-summary features were trained and evaluated separately within each colony, and the held-out predictions were pooled across colonies for visualization. These sex classifications did not exceed chance in any colony (all Holm-adjusted *P* > 0.05; one-sided within-colony label-permutation tests, 1,000 permutations). Cells show the percentage of animals in each true class assigned to each predicted class; fractions indicate the corresponding number of animals relative to the true-class total. The pooled balanced accuracy was 55.7%; NS, not significant.

**Supplementary Figure S4.**
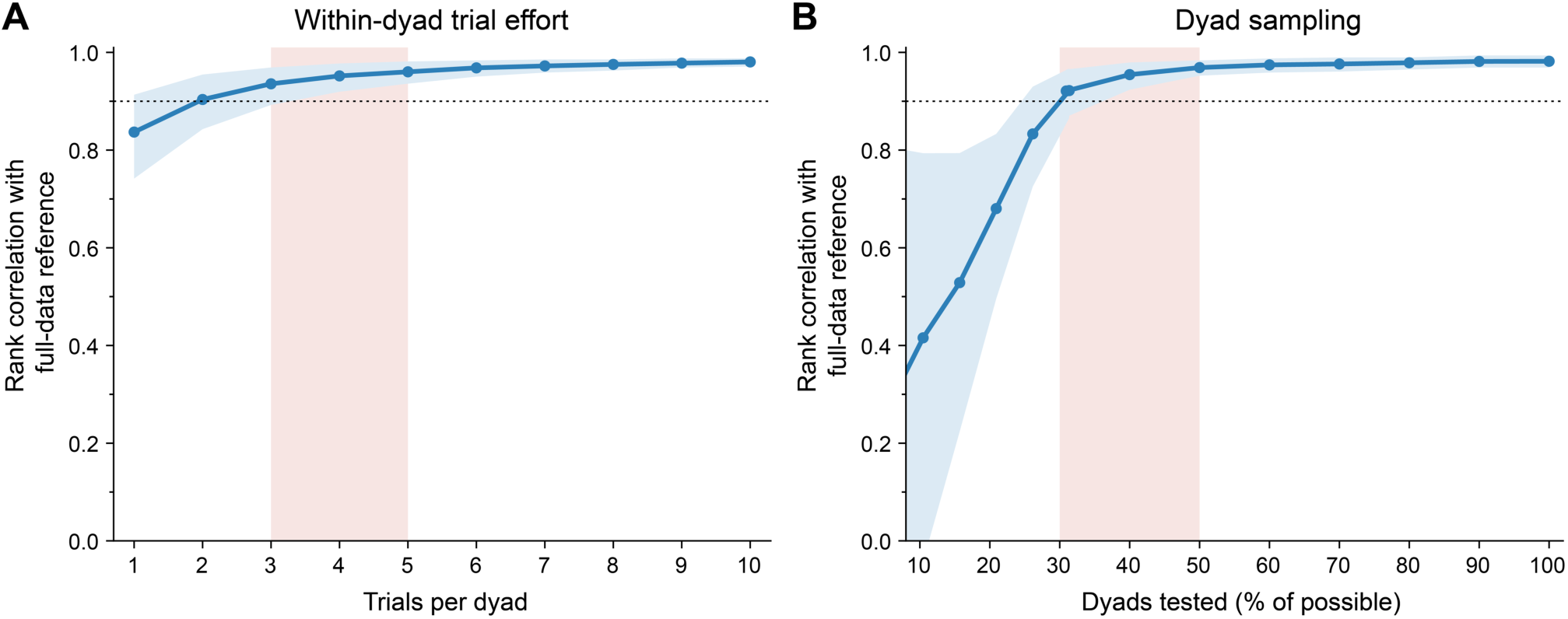
Sparse adaptive sampling efficiently recovers colony-wide dominance rank. Exhaustive, repeated dyad testing to rank animals becomes prohibitive as colony size increases. We therefore asked whether colony-wide rank could be recovered without compromising accuracy by testing fewer dyads and fewer trials per dyad. We first established a full-data reference by exhaustively testing colony B, completing 10 trials for all 153 possible dyads among 18 animals. We then retrospectively subsampled this dataset and compared the inferred ranks with the full-data rank order. (**A**) Rank correlation with the full-data reference at 40% dyad coverage when each dyad was tested for a fixed 1–10 trials. These results motivated a stopping rule in which testing ended after a 3–0 outcome but otherwise continued to five trials. (**B**) Rank correlation with the full-data reference as a function of the proportion of possible dyads using this stopping rule. In both panels, blue lines and points indicate means, and blue shading denotes the 10th–90th percentiles across 1,000 simulations per condition. Pale red bands indicate the selected operating ranges of 3–5 trials per dyad and 30%–50% dyad coverage, which we subsequently used to rank colonies H, P, and S. Black dotted lines indicate a rank correlation of 0.9.

**Supplementary Figure S5.**
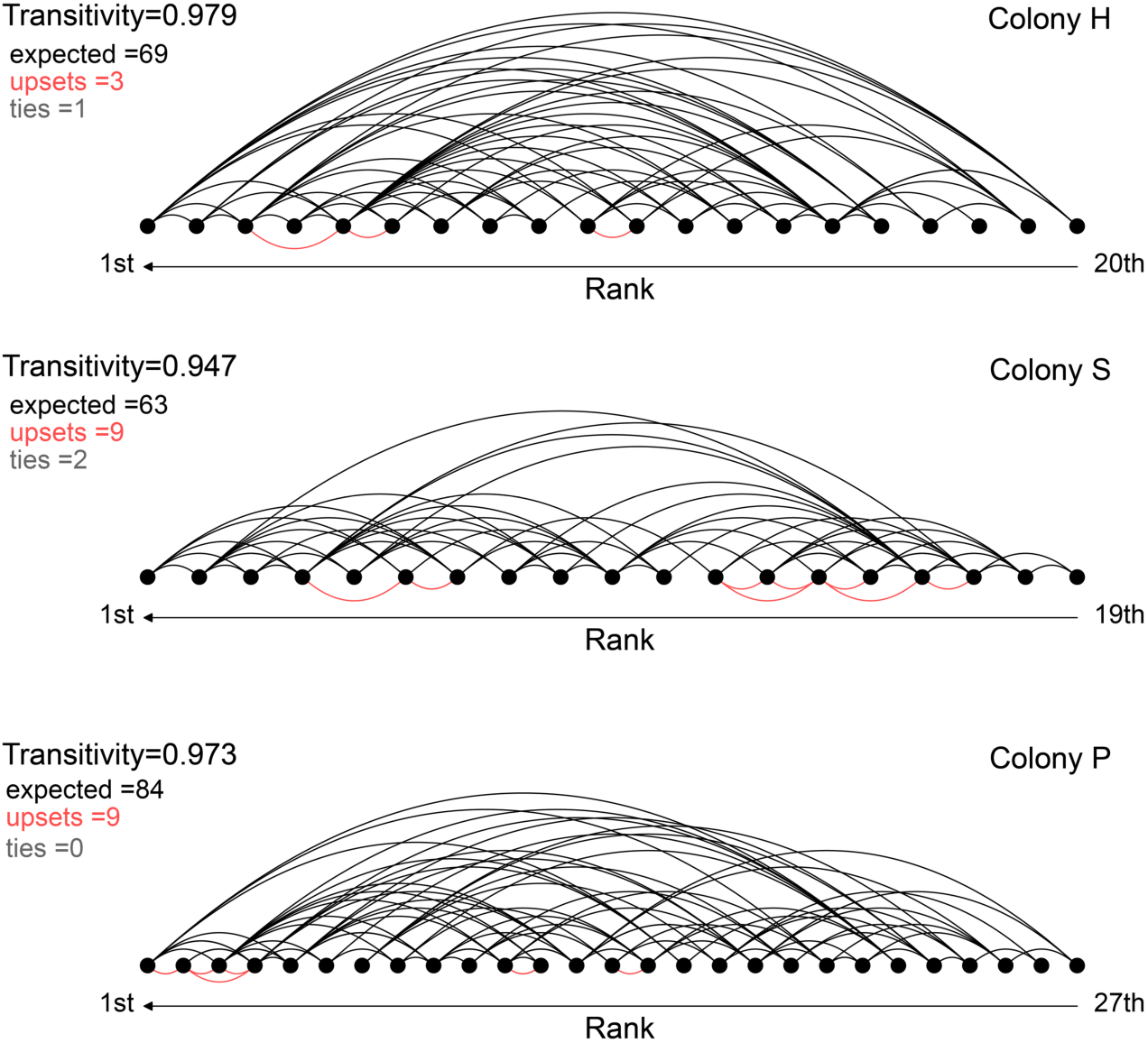
Tube-test dominance hierarchies are highly transitive across colonies, related to Figure 3. Rank-ordered dominance networks are shown for colony H (top), colony S (middle), and colony P (bottom). Each node represents an animal, ordered from highest to lowest rank from left to right. Black arcs indicate expected outcomes in which the higher-ranked animal won the dyad, whereas red arcs indicate upsets. Tied dyads are not connected. All three colonies exhibited highly transitive hierarchy structures (*P* < 0.001 for all colonies by permutation tests; see Methods).

**Supplementary Figure S6.**
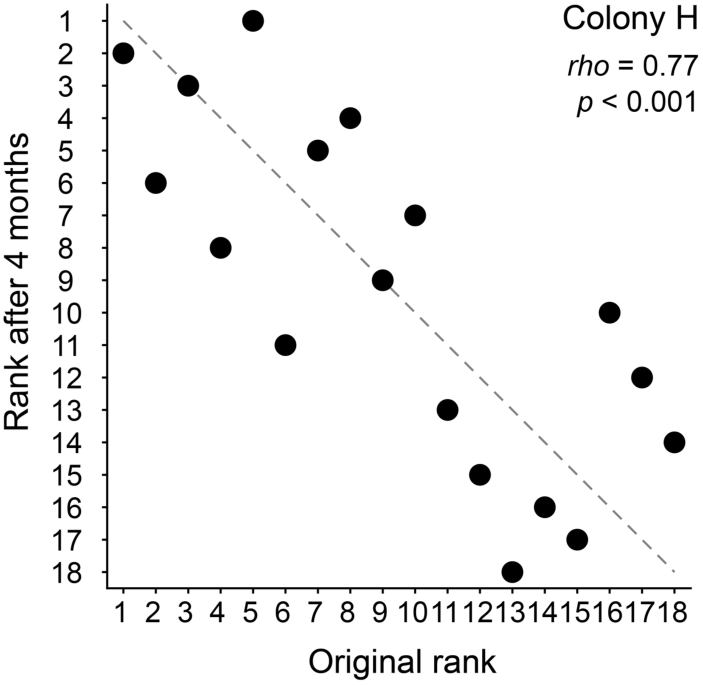
Rank remained stable over 4 months in colony. **H.** *rho* and *P*, Spearman’s rank correlation test.

**Supplementary Figure S7.**
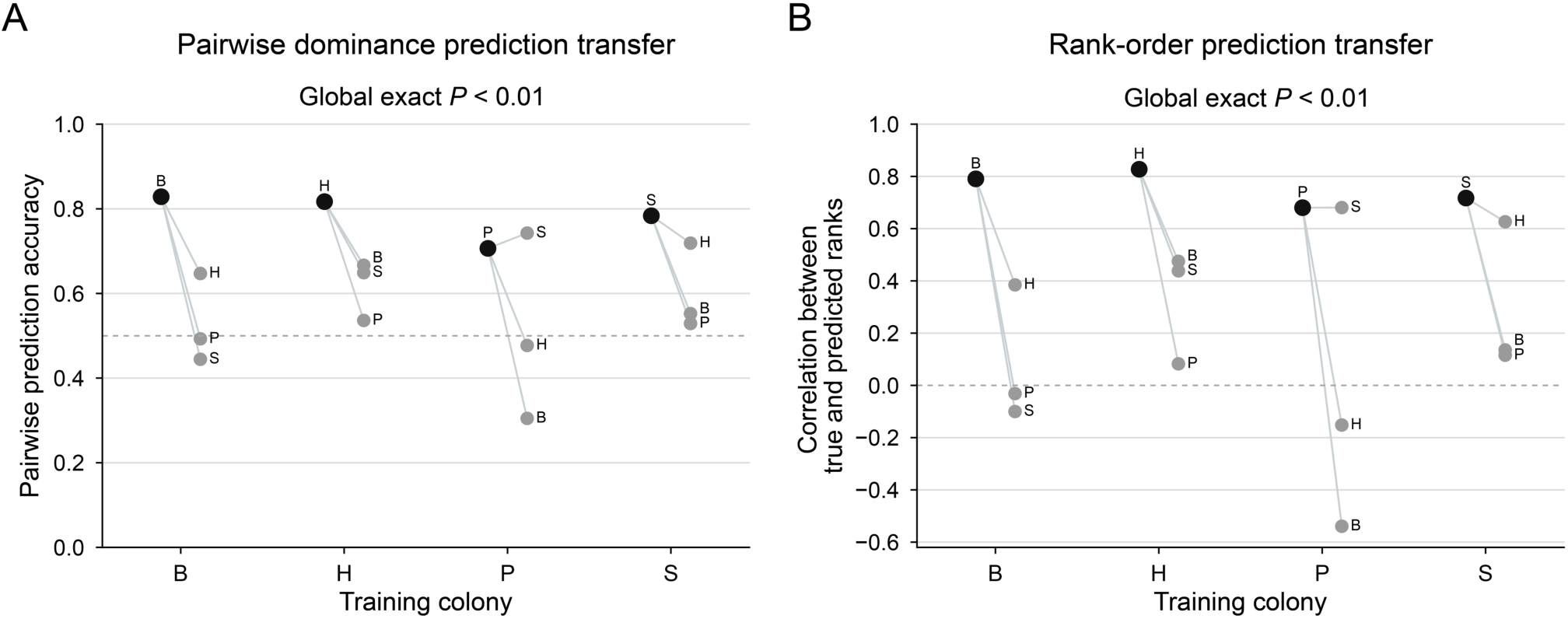
Dominance and rank prediction models transfer poorly across colonies. (**A**) Pairwise dominance-prediction accuracy of PLS regression models fitted in each colony and evaluated within the same colony (black) or transferred to each of the other three colonies (gray). (**B**) Correlations between true and predicted rank orders. Letters identify the target colony, and lines connect predictions made by the same training-colony model. Same-colony performance exceeded cross-colony performance for both pairwise dominance prediction (mean accuracy, 0.78 versus 0.56; *P* = 0.008) and rank-order prediction (mean Spearman’s *rho*, 0.75 versus 0.18; *P* = 0.008). *P* values were calculated using two-sided exact source-stratified permutation tests. Dashed lines indicate chance accuracy in A (50%) and zero correlation in B.

## Notes

### Competing Interest Statement

The authors have declared no competing interest.

